# Measuring hidden phenotype: Quantifying the shape of barley seeds using the Euler Characteristic Transform

**DOI:** 10.1101/2021.03.27.437348

**Authors:** Erik J. Amézquita, Michelle Y. Quigley, Tim Ophelders, Jacob B. Landis, Daniel Koenig, Elizabeth Munch, Daniel H. Chitwood

**Affiliations:** Department of Computational Mathematics, Science & Engineering, Michigan State University, East Lansing, MI, USA; Department of Horticulture, Michigan State University, East Lansing, MI, USA; Department of Mathematics, Michigan State University, East Lansing, MI, USA; Department of Mathematics and Computer Science, TU Eindhoven, Eindhoven, The Netherlands; Department of Information and Computing Sciences, Utrecht University, Utrecht, The Netherlands; School of Integrative Plant Science, Section of Plant Biology and the L.H. Bailey Hortorium, Cornell University, Ithaca, NY, USA; BTI Computational Biology Center, Boyce Thompson Institute, Ithaca, NY, USA; Department of Botany & Plant Sciences, University of California, Riverside, CA, USA

**Keywords:** Topological Data Analysis, Euler characteristic transform, mathematical biology, data science, shape

## Abstract

Shape plays a fundamental role in biology. Traditional phenotypic analysis methods measure some features but fail to measure the information embedded in shape comprehensively. To extract, compare, and analyze this information embedded in a robust and concise way, we turn to Topological Data Analysis (TDA), specifically the Euler Characteristic Transform. TDA measures shape comprehensively using mathematical representations based on algebraic topology features. To study its use, we compute both traditional and topological shape descriptors to quantify the morphology of 3121 barley seeds scanned with X-ray Computed Tomography (CT) technology at 127 micron resolution. The Euler Characteristic Transform measures shape by analyzing topological features of an object at thresholds across a number of directional axes. A Kruskal-Wallis analysis of the information encoded by the topological signature reveals that the Euler Characteristic Transform picks up successfully the shape of the crease and bottom of the seeds. Moreover, while traditional shape descriptors can cluster the seeds based on their accession, topological shape descriptors can cluster them further based on their panicle. We then successfully train a support vector machine (SVM) to classify 28 different accessions of barley based exclusively on the shape of their grains. We observe that combining both traditional and topological descriptors classifies barley seeds better than using just traditional descriptors alone. This improvement suggests that TDA is thus a powerful complement to traditional morphometrics to comprehensively describe a multitude of “hidden” shape nuances which are otherwise not detected.

## Introduction

There is a discrepancy between the information embedded in biological forms that we can discern with our senses versus that which we can quantify. Methods to comprehensively quantify phenotype are not commensurate with the thoroughness and speed with which genomes can be sequenced. High-throughput phenotyping has enabled us to collect large amounts of phenotyping data (Andrade-Sanchez et al., 2013; Araus and Cairns, 2014; Tanabata et al., 2012); nonetheless, we are not maximizing the information extracted from the data we collect.

One framework for extracting information embedded within data is to consider its shape. From a morphological perspective, the form of biological organisms is both data and literal shape simultaneously. Landmark-based approaches based on Procrustean superimposition (Bookstein, 1997) and Fourier-based decomposition of closed outlines (Kuhl and Giardina, 1982; Lestrel, 1997) comprise traditional morphometric methods. These approaches measure shape comprehensively, but are limited to either a geometric perspective that only considers the distances and relative positions of data points to each other or to a frequency domain transform of a closed contour. We thus turn to topology, the mathematical discipline that studies shape in a more abstract sense.

Topological Data Analysis (TDA) is a set of tools that arise from the perspective that all data has shape and that shape is data (Amézquita et al., 2020; Lum et al., 2013; Munch, 2017). TDA treats the data as if made of elementary building blocks: points, edges, squares, and cubes, referred to as 0−, 1−, 2−, and 3-dimensional *cells* respectively (Fig. 1A). A collection of cells is referred to as a *cubical complex*, or complex, for short.

**Figure 1:**
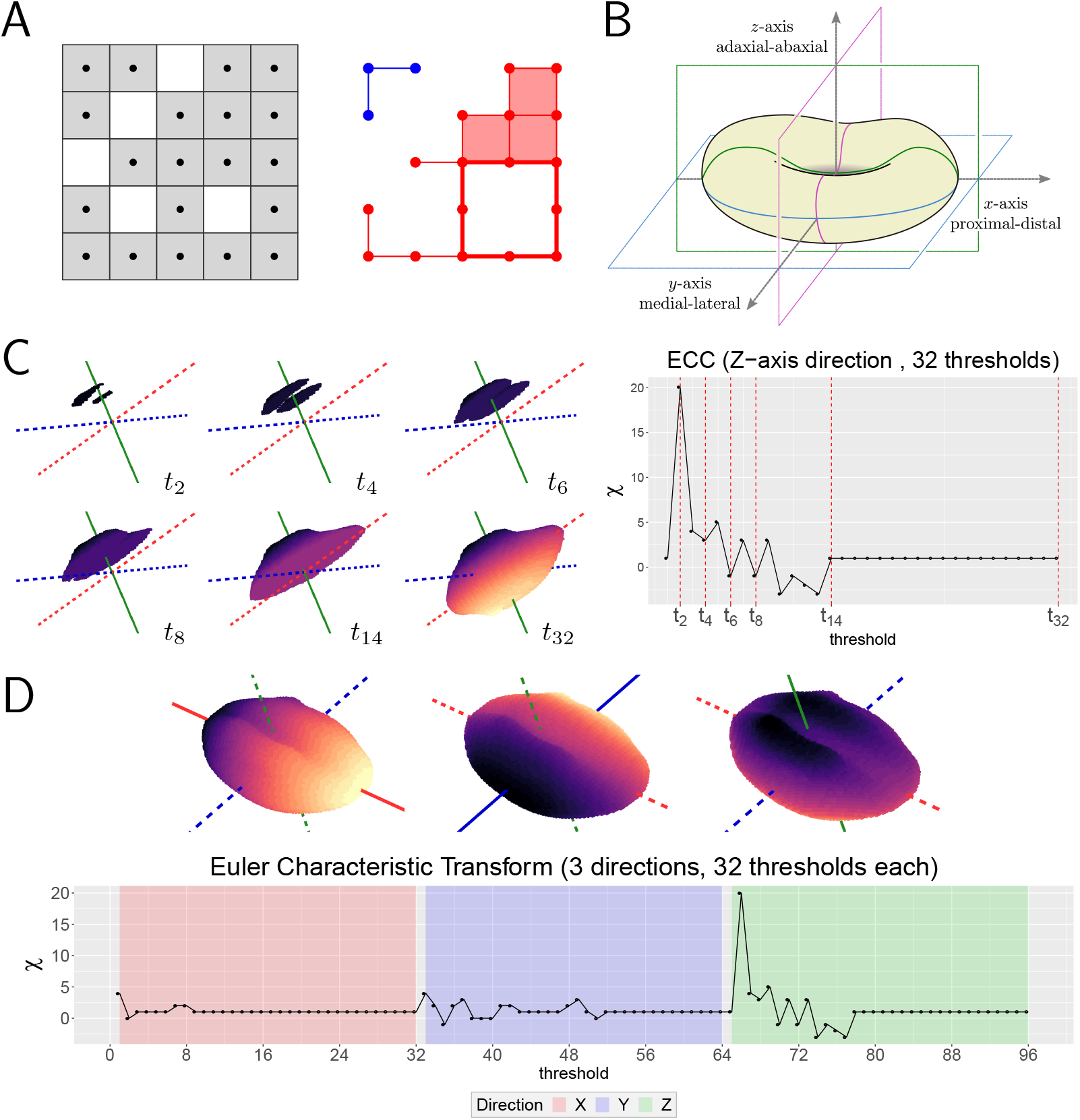
Extracting topological shape signatures from barley seeds. **A.** A binary image (left) is treated as a cubical complex (right). This cubical complex has 2 connected components, 1 loop, 0 voids. The distinct connected components are colored in blue and red respectively. The loop is emphasized with thicker edges. **B.** The barley seeds were aligned so that their proximal-distal, medial-lateral, and adaxial-abaxial axes corresponds to the *X, Y, Z*-axes in space. **C.** Example of an Euler Characteristic Curve (ECC) as we filter the barley seed across the adaxial-abaxial axis (depicted as a solid, green line) through 32 equispaced thresholds. **D.** The Euler Characteristic Transform (ECT) consists of concatenating all the ECCs corresponding to all possible directions. In this example, we concatenate 3 ECCs corresponding to the *X, Y, Z* directions, represented by the solid lines respectively.

Cubical complexes are both a natural and consistent way to represent image data (Kovalevsky, 1989). Given a grayscale image, we follow a strategy similar to Wagner et al. (2012) to construct a cubical complex: a nonzero pixel will correspond to a vertex in our complex. If two pixels are adjacent —in the 4-neighborhood sense— we say that there is an edge between the corresponding vertices in the complex. If 4 pixels in the image form a 2 × 2 square, we will consider a square in our complex between the corresponding 4 vertices (Fig. 1A). Additionally, for the 3D image case, if 8 voxels—the 3D equivalent of pixels—make a 2 × 2 × 2 cube, we will draw a cube in our complex between the corresponding 8 vertices.

TDA seeks to describe the shape of our data based on the number of relevant topological features found in the corresponding complex. For instance, the complex in Figure 1A has two distinct, separate pieces colored in blue and red respectively, formally referred to as *connected components*. This complex also has 8 edges forming the outline of a square without an actual red block filling it—edges thickened for emphasis—this is referred to as a *loop*. In higher dimensions, we could also consider hollow blocks containing *voids*. We can even go a step further and summarize these topological features with a single value known as the *Euler characteristic*, represented by the Greek letter *χ*, defined for voxel-based images as

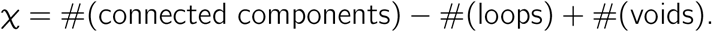

The Euler characteristic is a topological invariant; that is, it will remain unchanged under any smooth transformation applied to our shape. The well-known but surprising Euler-Poincaré formula states that *χ* can be computed easily as

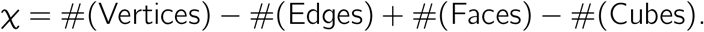

This equivalence can be seen in the cubical complex in Figure 1A, where

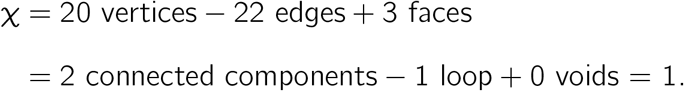

The Euler characteristic by itself might be too simple. Nonetheless, we can extract more information out of our data-based complex if we think of it as a dynamic object that grows in number of vertices, edges, and faces across time. As our complex grows, we may observe significant changes in *χ*. The changes in *χ* can be thought as a topological signature of the shape, referred to as an *Euler characteristic curve (ECC)*. The growth of the complex is defined by a *filter function* which assigns a real number value to each voxel. For reasons discussed later, we will focus on directional filters which assign to each voxel its height as if measured from a fixed direction.

As an example, consider the cubical complex of a barley seed and the direction corresponding to the adaxial-abaxial axis (Fig. 1B). Voxels at the top of the seed will be assigned the lowest values, while voxels at the bottom will obtain the highest values. We then consider 32 equispaced, increasing thresholds *t*_1_ < *t*_2_ < … < *t*_32_ which define 32 different slices of equal thickness along the adaxial-abaxial axis. We start by computing the Euler characteristic of the first slice, that is, all the voxels with filter value less than *t*_1_. Next we aggregate the second slice, which are all the voxels with filter value less than *t*_2_, and recompute the Euler characteristic. We repeat the procedure for the 32 slices. For instance in Figure 1C, we observe that we started with scattered voxels which are thought of as many connected components which may explain the high Euler characteristic values. As we keep adding slices, we connect most of the stray voxels into fewer but larger connected components, and simultaneously, we might have created loops as seen in *t*_4_ and *t*_6_. This merging of connected components, and formation and closing of loops might explain the fluctuation of the Euler characteristic between positive and negative values. Finally, after more than half of the slices have been considered, at *t*_14_, we observe that no new loops are formed, and every new voxel will simply be part of the single connected component. Thus, the Euler characteristic remains constant at 1. The ECC is precisely the sequence of different Euler characteristic values as we add systematically individual slices along the chosen direction.

To get a better sense of how the Euler characteristic changes overall, we can compute several ECCs corresponding to different directional filters. For example, in Figure 1D we choose three directions in total corresponding to the proximal-distal, medial-lateral, and adaxial-abaxial axes respectively. Each filter produces an individual ECC, which we later concatenate into a unique large signal known as the *Euler Characteristic Transform (ECT).*

There are two important reasons to use ECT over other TDA techniques. First, the ECT is computationally inexpensive, since it is based on successive computations of the Euler characteristic, which is simply an alternating sum of counts of cells. This inexpensiveness is especially relevant as we are dealing with thousands of extremely high-resolution 3D images. Assuming that we have already treated the image as a cubical complex, we can compute a single ECC in linear time with respect to the number of voxels in the image (Richardson and Werman, 2014). We can thus compute the ECT of a 50,000-voxel seed scan with 150 directions in less than two seconds on a traditional PC. The second reason to use the ECT is its provable invertibility and statistical sufficiency. As proved by Turner et al. (2014), and later extended separately by Curry et al. (2018) and Ghrist et al. (2018), if we compute all possible directional filters we would have sufficient information to reconstruct the original shape. Moreover, this ECT is a sufficient statistic that effectively summarizes all information regarding shape. Although there are infinite possible directional filters, there is ongoing research into defining a sufficient finite number of directions such that we can effectively reconstruct shapes based solely on their finite ECT (Belton et al., 2020; Betthauser, 2018; Curry et al., 2018; Fasy et al., 2019). Nonetheless, a computationally efficient reconstruction procedure for large 3D images remains elusive.

Another computational consideration is the fact that the ECT produces a vector of topological information of #(directions) × #(thresholds) dimensions, which is usually above 2000 dimensions. In general, high-dimensional vectors tend to produce distorted prediction and regression results (Köppen, 2000), and it is advised to denoise and summarize these vectors by using different dimension reduction techniques. One such standard technique is principal component analysis (PCA), which seeks to project the high-dimensional vectors unto the orthogonal directions that capture the greatest variability of the data. These linear directions are referred to as the *principal components* of the data. Sometimes, the data cannot be properly summarized as a collection of lines. A more flexible approach is to consider *kernel PCA (KPCA)* (Schölkopf et al., 1998), an *nonlinear* alternative. By specifying a *kernel function*, we can instead project the high-dimensional samples unto the polynomial, trigonometric, or circular curves that capture the most variance of the data. A completely different dimension reduction strategy is the *uniform manifold approximation and projection (UMAP)* (McInnes et al., 2020), which also draws several ideas from TDA. Intuitively, UMAP seeks to project the high-dimensional data unto a low-dimensional space while preserving the most prominent topological local features. That is, if the original data contains large connected components, wide loops, and ample voids, its low-dimensional UMAP projection should also exhibit several connected components, loops, and voids. If two sample points are in the same connected component in the high-dimensional space, these two should remain in the same cluster when projected to the low-dimensional space.

Here we show the use of ECTs to correctly summarizes the shape of barley seeds as a proof of concept. We scanned a collection of barley panicles comprising 28 different accessions with X-ray CT technology at 127 micron resolution. These scans were later digitally processed to isolate 3121 individual grains, and their morphology was quantified using both traditional and topological shape descriptors. We then explored both qualitatively and quantitatively the descriptiveness of these measurements. To aid both assessments, we used KPCA and UMAP separately to aggressively reduce the dimension of the traditional and ECT vectors. We observe that traditional shape descriptors tend to cluster seeds based on their accession, while KPCA-reduced topological shape descriptors tend to cluster them based on panicles. UMAP-reduced topological descriptors balance both approaches and draw shape distinctions at both accession and spike level. This in turn shows that KPCA and UMAP draw from different pieces of ECT information. This observation suggests that the ECT effectively summarizes both spike-specific and accession-specific morphological information which can be then highlighted with an appropriate dimension reduction technique. To quantify the descriptor correctness, we trained a support vector machine (SVM) to determine the accession of individual grains based on their shape alone. Our experiments show that SVMs perform better whenever topological information is taken into account, which suggests that the ECT measures shape that is “hidden” from traditional shape descriptors.

## Materials and Methods

We selected 28 barley accessions with diverse spike morphologies and geographical origins for our analysis (Harlan and Martini, 1929, 1936, 1940). In November of 2016, seeds from each accession were stratified at 4C on wet paper towels for a week, and germinated on the bench at room temperature. Four day old seedlings were transferred into pots in triplicate and arranged in a completely randomized design in a greenhouse. Day length was extended throughout the experiment using artificial lighting —minimum 16h light / 8h dark. After the plants reached maturity and dried, a single spike was collected from each replicate for scanning at Michigan State University. The scans were produced using the North Star Imaging X3000 system and the included efX software, with 720 projections per scan, with 3 frames averaged per projection. The data was obtained in continuous mode. The X-ray source was set to a voltage of 75 kV, current of 100 *μ*A, and focal spot size of 7.5*μ*m. The 3D reconstruction of the spikes was computed with the efX-CT software, obtaining a final voxel size of 127 microns. The intensity values for all raw reconstructions was standardized as a first step to guarantee that the air and the barley material had the same density values across all scans. Next, the air and debris was thresholded out, and awns digitally pruned (Figs. 2A-D). Finally, the seed coat of the caryopses was digitally removed, leaving only the embryo and endosperm due to their high water content (Fig. 2E). We did not have enough resolution in the raw scans to distinguish clearly the endosperm from the embryo. Hereafter, we will refer to these embryo-endosperm unions simply as seeds. Thus, we digitally isolated all the seeds and obtained a collection of 3438 seeds in total. Due to the large volume of data, we used an in-house scipy-based python script to automate the image processing pipeline for all panicles and grains.

**Figure 2:**
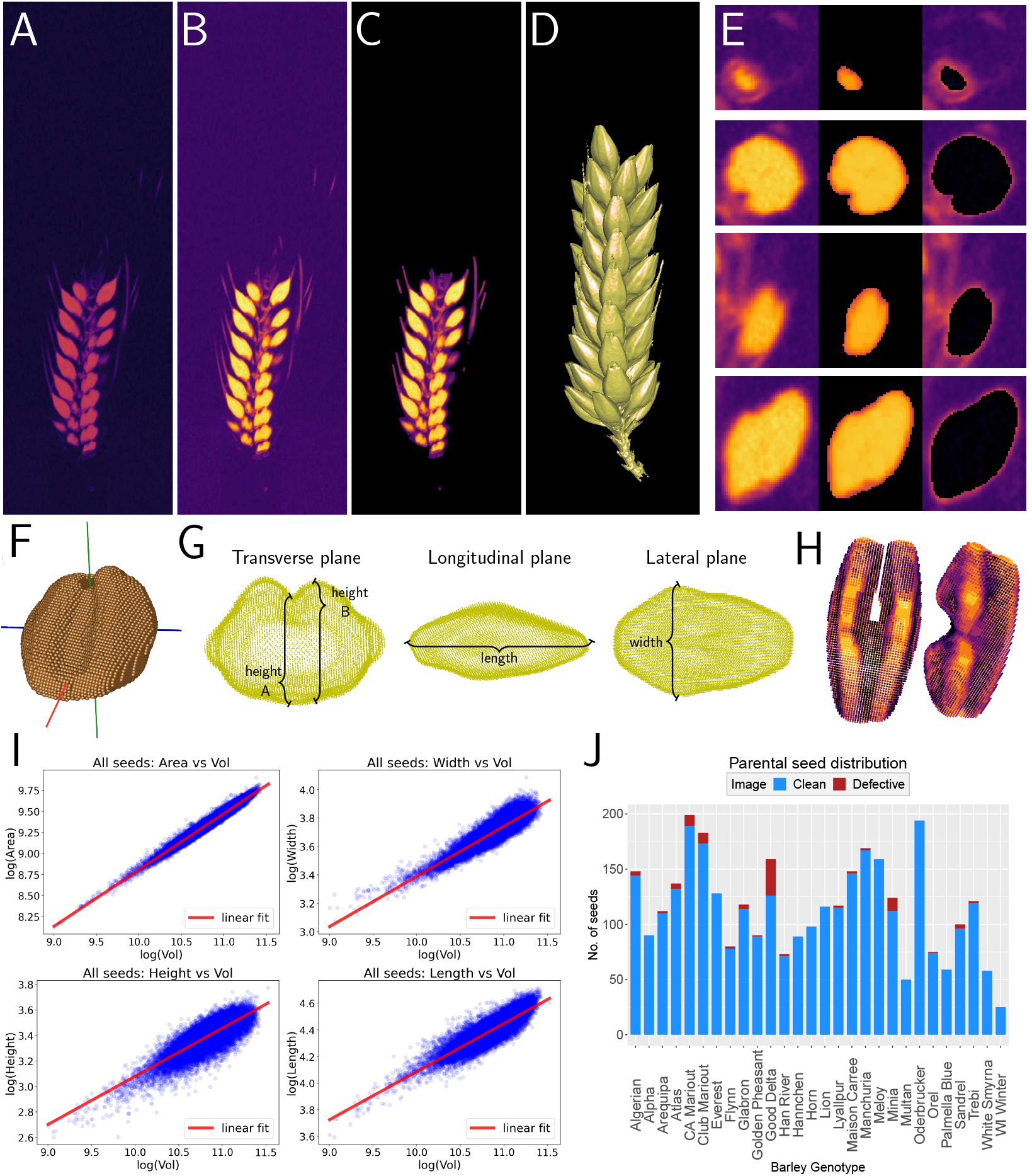
Barley image processing. The morphology measurements were extracted from 3D voxel-based images of the barley panicles. Before any analysis was done, the **A.** raw X-ray CT scans of the panicles had their **B.** densities normalized, **C.** air and other debris removed, and awns pruned. **D.** After automating these image processing steps, we could finally work with a large collection of clean, 3D panicles. **E.** An extra digital step segmented the individual seeds (embryo and endosperm) for each barley spike. The left shows the original raw scan, the center shows the isolated seed, while the right side shows part of the coat that was removed while segmenting. **F.** The seeds were aligned according to their principal components, which allowed us to **G.** measure a number of traditional shape descriptors. **H.** The incomplete or broken seeds were **I.** initially identified as out-liers of the allometry plots. These damaged seeds were later removed from the data set. **J.** The total number of clean and defective seeds measured from each accession. Defective seeds were not concentrated in a particular accession.

To make the collection of different directional filters comparable across seeds, all the seeds were aligned with respect to their first three principal components. Since all the seeds are oblong in shape, this PCA-based alignment corresponds to the proximal-distal, medial-lateral, and adaxial-abaxial axes respectively (Fig. 1B; Fig. 2F). The orientation of the principal components is arbitrary with every run, so we did keep track of the crease and the tip of seed and flipped the axes accordingly so that the tip would always be located as the rightmost point of the image and the crease would always point north. With this uniform alignment we were able to measure the length, width, heights, surface area and volume of each seed (Fig. 2G). We also computed the convex hull for each seed and measured its surface area and volume, as well as the ratios with respect to seed surface area and volume. In total, 11 different traditional shape descriptors were measured. Damaged and incomplete seeds (Fig. 2H) were removed by evaluating allometry plots along their best linear fits and residuals (Fig. 2I). Points with residuals 3 times larger than the standard deviation were deemed as outliers and the associated seed removed. A final visual assessment of the remaining images was conducted to ensure the removal of all damaged seeds. These outliers did not represent a significant portion of the seeds of any accession (Fig. 2J). In total we obtained 3121 cleanly segmented seeds. Every accession is represented on average by 111 seeds, with *±*42 seeds as standard deviation. All the accession numbers are within 2 standard deviations from this empirical mean (Fig. S1; Table 1).

**Table 1:**
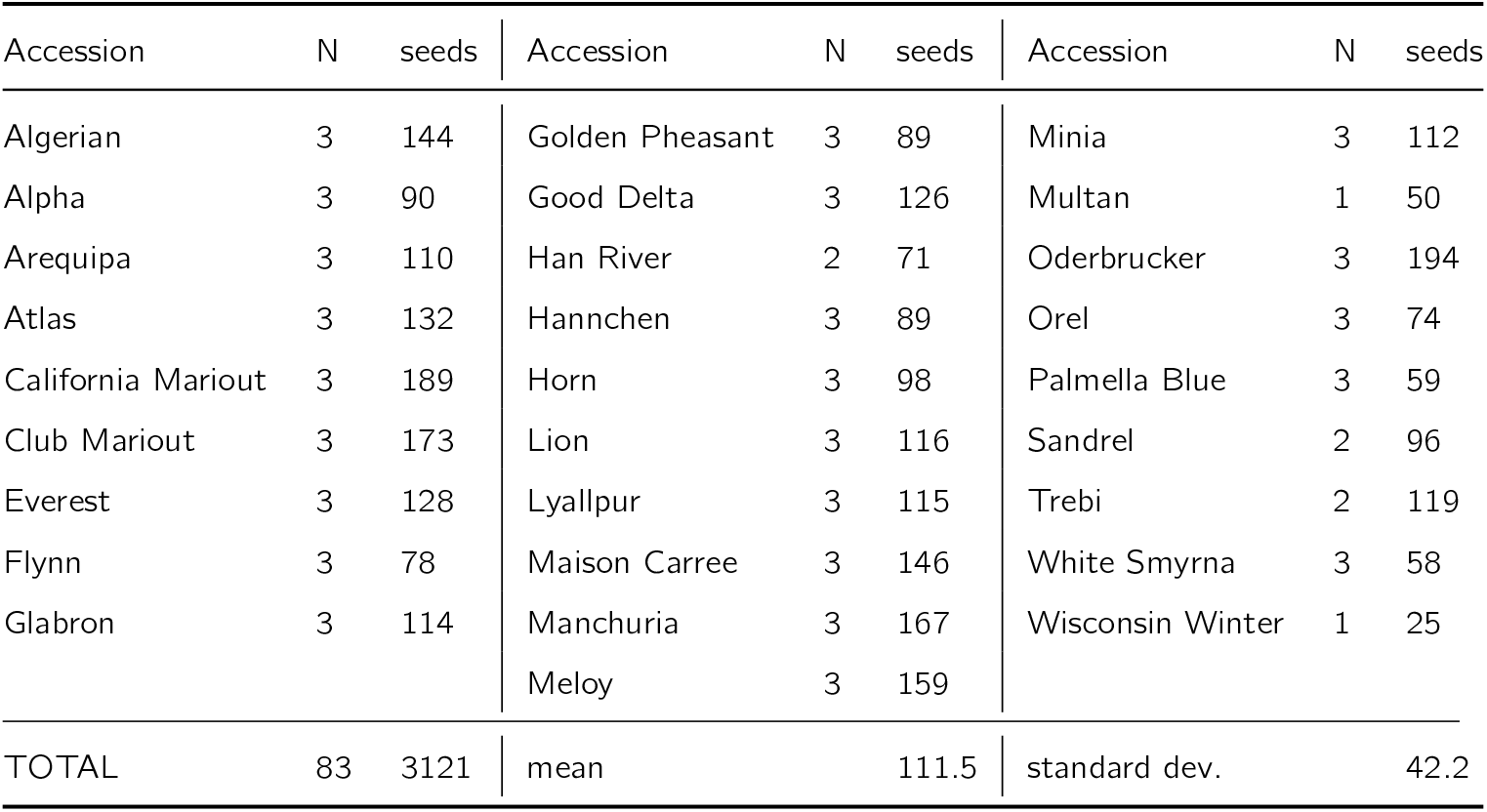
Sample size of seed scans used for each individual accession. **N** equals the number of panicles from which seeds are derived.

As a proof of concept, we explored how topological descriptors varied as we varied both the number of different directions and the number of uniformly spaced thresholds. In total, for every seed we computed the ECT considering 74, 101, 158, and 230 different directions. We emphasized directions toward the seed’s crease, which correspond to directions close to both north and south poles (Fig. 1B; Fig. 3). For each direction, we produced ECCs with 4, 8, 16, 32, and 64 thresholds.

**Figure 3:**
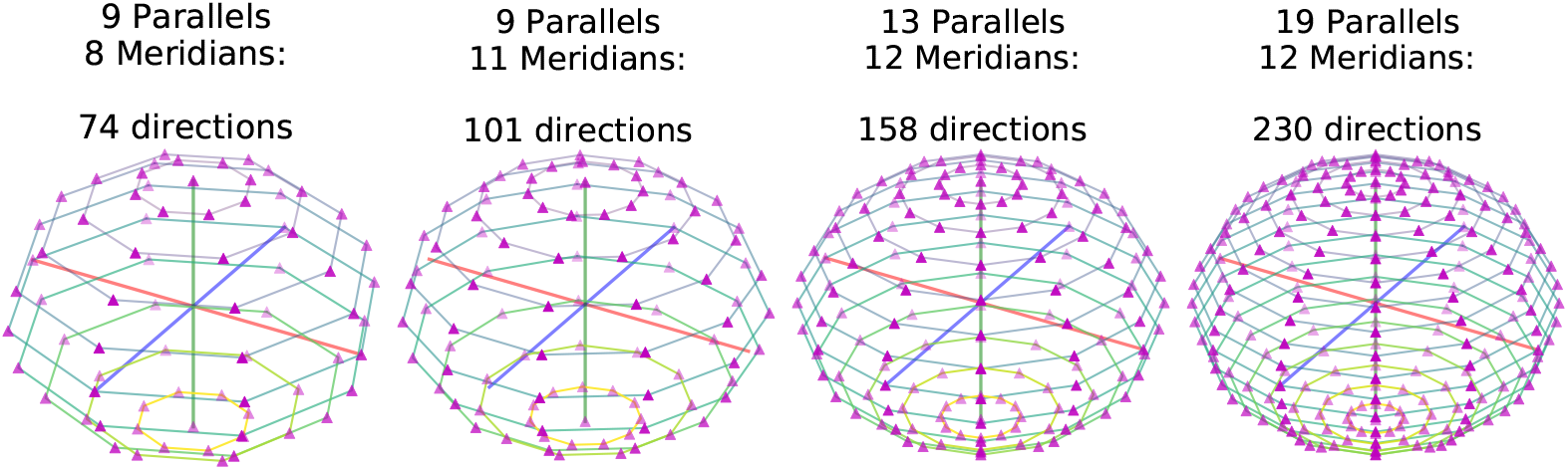
Directions chosen to compute the ECT. The sphere was split into a equispaced fixed number of parallels and meridians in each case. The directions were the taken from the intersections.

Recall that the ECT is a record of how topology changes at every single slice taken at every direction (Fig. 1C). We performed Kruskal-Wallis one-way analyses (Kruskal and Wallis, 1952) to determine if the Euler characteristic interaccession variance was significantly different than the intra-accession variance at a particular slice and direction. This way, we observed which parts of the seed anatomy were of particular relevance to the ECT. Accessions and individual spikes were both considered as possible classes when performing the Kruskal-Wallis tests. These results follow a conservative 10^−10^ false discovery rate after considering a multiple test Benajamini-Hochberg correction (Benjamini and Hochberg, 1995).

For every seed we computed a very high-dimensional vector of topological information, usually above 2000 dimensions, which were later reduced in dimension independently with KPCA and UMAP to prevent high-dimensionality distortions. A non-linear KPCA with a *σ* = 1 Laplacian kernel reduced the ECT dimension based on its largest source of variance. UMAP on the other hand was used to preserve the prominent, high-dimensional topological features of the ECT in an unsupervised fashion. We fixed the use of 50 nearest neighbors, 0.1 minimum distance, and Manhattan distance as the rest of key UMAP hyperparameters. For all dimension reduction techniques, the ECT dimension was reduced to just 2, 3, 6, 12, and 24 dimensions. We focused on an aggressive 2-dimensional reduction for visualization purposes both with KPCA and UMAP.

To evaluate the descriptiveness, we trained three non-linear support vector machines (SVM) with radial kernel *σ* = 0.1 (Burges, 1998) to characterize and predict the seeds from 28 different accessions based on three different collections of descriptors: traditional, topological, and combining both traditional and topological descriptors. In every case, the descriptors were centered and scaled to variance 1 prior to classification. Given that SVM is a supervised learning method, we partitioned our data into training and testing sets. In our case, we randomly sampled 75% of the seeds from every accession as our training data set, labeled according to their accession. The remaining 25% was used to test the accuracy of our prediction model. We repeated this SVM setup 100 times and considered the average accuracy and confusion matrices as final results. This was done for all possible combinations of directions, thresholds, and dimensionality reductions mentioned above. The SVM was our classifier of choice since it is quick to train and it does not require vast amounts of training data to produce reasonable results.

## Results

Topological and combined shape descriptors tend to produce more accurate shape-based classification results, provided that the ECT is computed with sensible parameters and an adequate dimension reduction technique. The best SVM classification results were yielded by topological and combined shape descriptors based off a 2568-dimensional ECT —158 directions and 16 thresholds (Fig. S3). Based on the highest *F*_1_ classification scores, these high-dimensional vectors were best parsed after being reduced to just 2 dimensions with KPCA, or to 12 dimensions with UMAP. Hereafter, the rest of topological related results are based on these specific choice of directions, thresholds, and dimensionality reduction.

A Kruskal-Wallis one-way analysis of the ECT vectors, combined with a Benjamini-Hochberg correction admitting a 10^−10^ FDR, reveals 55 features that explain the most of inter-accession variance (Fig. 4A). The most accession-discerning slices and directions correspond to the north and south poles (Fig. 4B). As discussed in the seed alignment heuristics in the Methods section, these pole directions in turn correspond to the morphology of the crease and the bottom of the seed (Fig. 4C). Similar results were observed when analyzing for the most spike-discerning directions (Fig. S4). In other words, the topological shape descriptors do measure the crease and bottom shape of the seed, a morphological feature not explicitly measured by our traditional setting.

**Figure 4:**
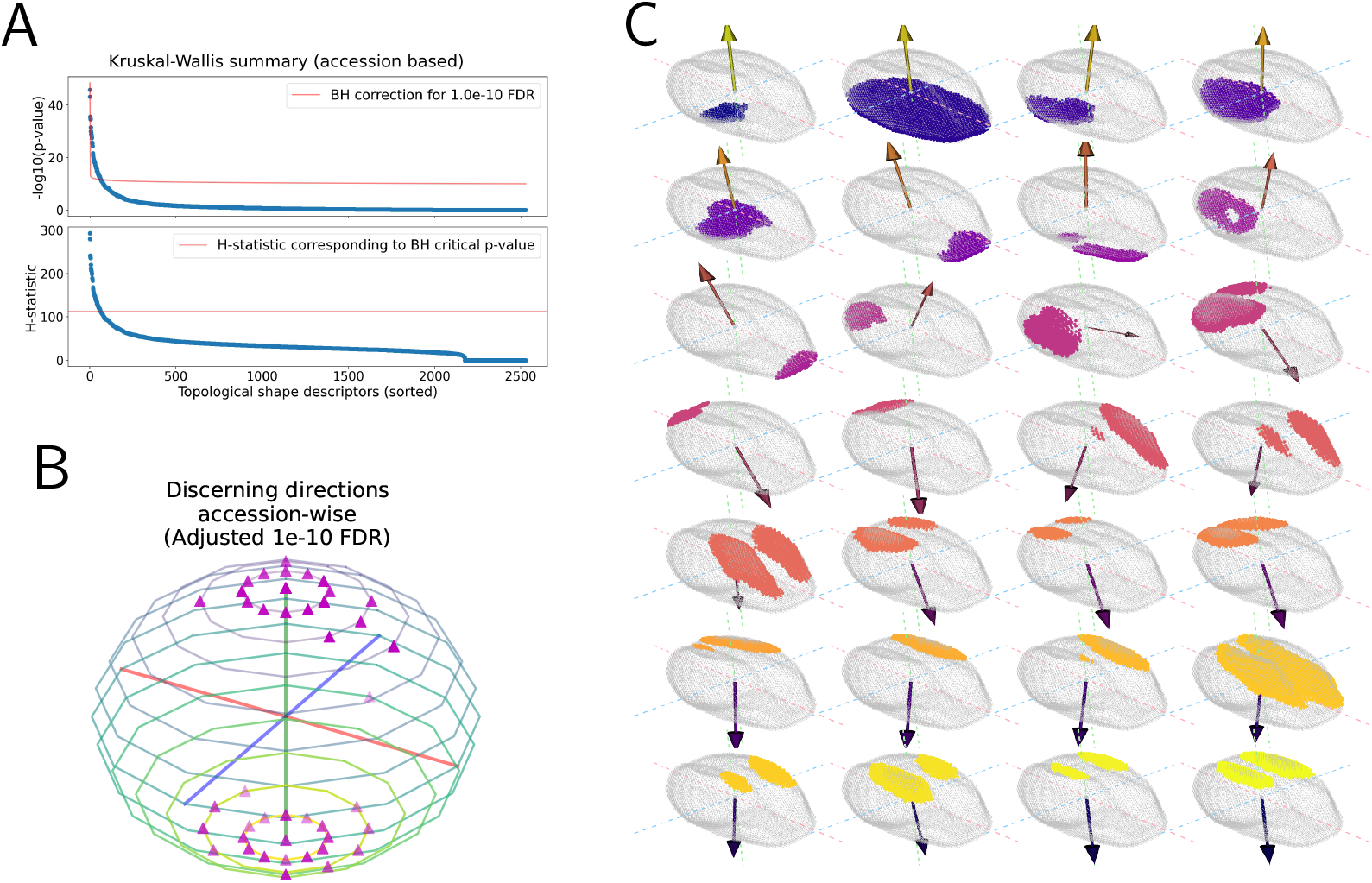
Relevant ECT directions and slices. **A.** We examine the inter-accession and intra-accession variance differences of the Euler characteristic for each direction and threshold. A Kruskal-Wallis analysis combined with a Benjamini-Hochberg multiple test correction suggests a handful of particularly discerning slices across accessions. **B.** These directions and thresholds are mostly concentrated around the poles, and **C.** correspond to the seed’s crease and bottom morphology.

Turning back to the traditional shape descriptors, these share similar distributions across the 28 accessions, provided they are all centered and scaled to variance 1 (Fig. 5A). Kruskal-Wallis analyses suggest that the seed length, surface area, and volume related measures explain the most inter-accession variance (Fig. S5A-B). Reducing the descriptors to a 2D representation with PCA suggests that these traditional descriptors tend to group the seeds based on their accession (Fig. 5B). These two components explain 84.0% of the total variance, with the first principal component explaining a considerable 72.2% alone. A similar grouping-by-accession behavior was observed whenever we reduced the traditional shape descriptors to 2 dimensions with UMAP instead. KPCA dimension reduction did not yield insightful results.

**Figure 5:**
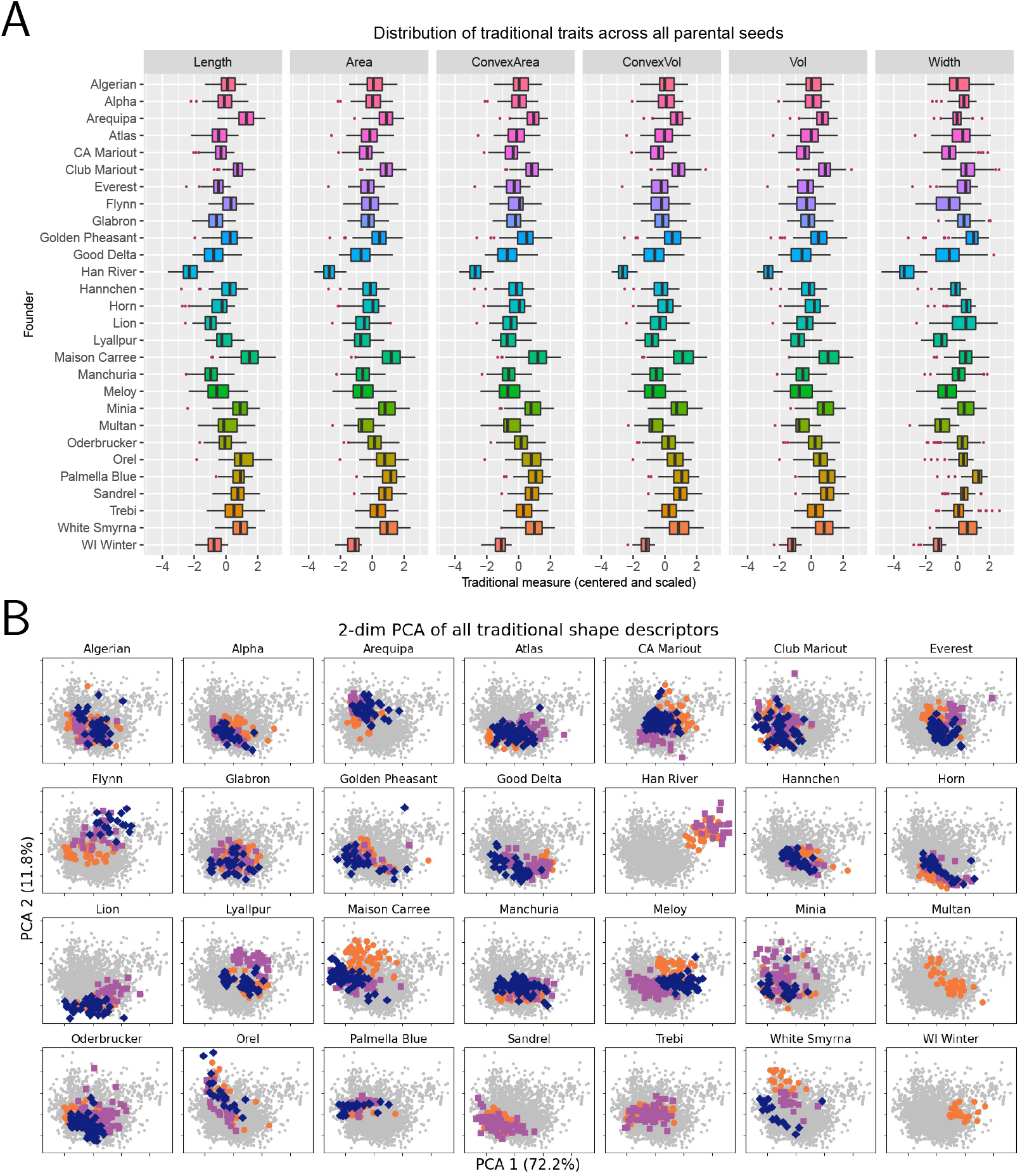
Distribution of traditional shape descriptors. **A.** Distribution of six of the 11 traditional seed shape descriptors across the 3121 seeds. These measurements were first centered at 0 and scaled to have variance 1. **B.** Plot of the first 2 principal components of the 11 shape descriptors. The first PC describes more than 70% of the total variance. Different marker and color indicate seeds from different spikes.

Topological shape descriptors on the other hand can provide a more spike-specific morphology encoding, depending on the dimension reduction technique used to parse the ECT. KPCA summarizes the topological information as a loop, with sharply defined clusters corresponding to seeds from individual spikes (Fig. 6A). On the other hand, the UMAP projection produces a large, round cluster. Notice that seeds of different spikes tend to lie on different locations, while these locations overlap partially for spikes of the same accession (Fig. 6B). This behavior suggests that UMAP dimension reduction tries to balance both spike-specific and accession-specific shape features.

**Figure 6:**
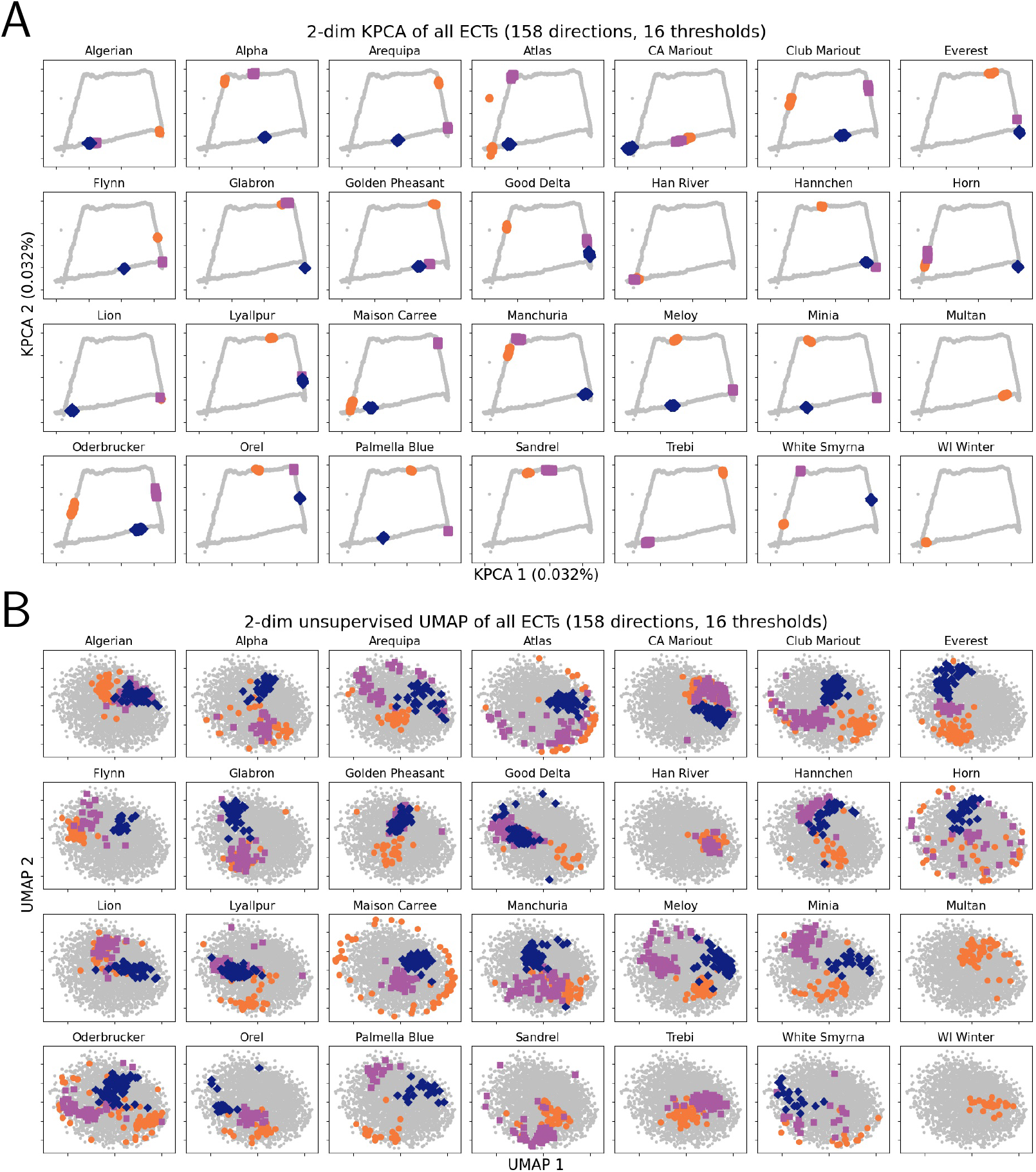
Dimension reduction of the ECT vectors. The ECT can produce a high-dimensional topological signature for each seed. To better visualize this topological information, we can reduce it to just two dimensions with **A.** kernel PCA or **B.** unsupervised UMAP. The seeds of individual accessions are highlighted in every frame. Different marker and color indicate seeds from different spikes.

Another round of Kruskal-Wallis analyses on the combined shape descriptors reinforce the idea that traditional descriptors cluster based on accession, KPCA-reduced topological descriptors do so based on spike, while UMAP-reduced ones provide a balanced clustering. The most inter-accession variance is explained predominantly by the traditional shape descriptors, with just a few topological features as complement (Figs. S5A-B). However, most of the inter-spike variance is predominantly captured by the dimension-reduced topological descriptors. The first two KPCA components do explain most of this inter-spike variance, which agrees with the tight panicle clusters seen before (Fig. 6; Fig. S5C). On the other hand, UMAP distributes regularly the spike variance across most of its components, complemented by a few traditional shape descriptors (Fig. S5D). In other words, traditional shape descriptors capture accession-specific shape features, KPCA highlights spike-specific features, and UMAP provides a balance between both of them.

When evaluating quantitatively the descriptiveness of these cluster differences, we observed that topological shape descriptors are able to produce much better SVM classification results than traditional shape descriptors (Table 2). Using exclusively traditional descriptors, the machine is able to correctly determine the grain variety roughly 57% of the time. For comparison, by simply randomly guessing the variety, we would expect to be correct just 1/28 × 100 ≈ 4% of the time. The classification could not be improved by reducing the dimension of the traditional vector (Fig. S2). If we use exclusively topological shape descriptors instead, the machine can classify different accessions with more than 75% accuracy. These results depend on the dimension reduction technique of choice (Fig. S3A). We observe that KPCA provides a powerful 2-dimensional summary of the ECT, which later can be used to predict grain accession with 85% classification accuracy. This accuracy diminishes considerably as more nonlinear principal components are considered. This drop of classification performance can be offset by combining the KPCA summary with traditional shape descriptors, which keep the classification accuracy above 70% (Fig. S3B).

**Table 2:**
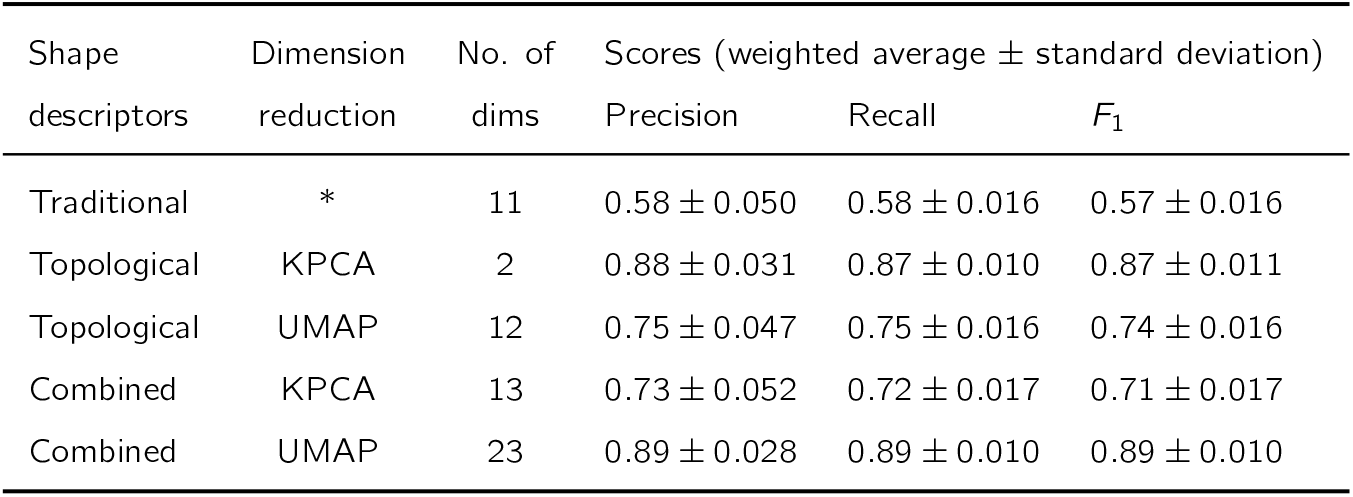
SVM classification accuracy of barley seeds from 28 different founding lines after 100 randomized training and testing sets. Since we are in a multi-class classification setting we first computed the precision, recall, and *F*_1_ scores for each founding line. Later, we computed the weighted average for each score, where the weight depended on the number of test seeds for each of the barley lines. Observe that the use of either topological or combined descriptors outperforms the use of exclusively traditional descriptors.

| Shape descriptors | Dimension reduction | No. of dims | Scores (weighted average ± standard deviation) | | |
| --- | --- | --- | --- | --- | --- |
| | | | Precision | Recall | $F_1$ |
| Traditional | * | 11 | $0.58 \pm 0.050$ | $0.58 \pm 0.016$ | $0.57 \pm 0.016$ |
| Topological | KPCA | 2 | $0.88 \pm 0.031$ | $0.87 \pm 0.010$ | $0.87 \pm 0.011$ |
| Topological | UMAP | 12 | $0.75 \pm 0.047$ | $0.75 \pm 0.016$ | $0.74 \pm 0.016$ |
| Combined | KPCA | 13 | $0.73 \pm 0.052$ | $0.72 \pm 0.017$ | $0.71 \pm 0.017$ |
| Combined | UMAP | 23 | $0.89 \pm 0.028$ | $0.89 \pm 0.010$ | $0.89 \pm 0.010$ |

The 2-dimensional UMAP summary (Fig. 6B) exhibits difficulties and discerning accessions, where classification accession does not go above 25%. Nonetheless, if a 12-dimension UMAP summary is considered, it is possible to classify accessions with 75% accuracy using exclusively topological information. Moreover, these UMAP-summary classification results can be further improved by combining them with traditional shape descriptors, where classification accuracy goes beyond 88%. The ECT thus captures important morphological patterns that can be complemented by size features which are provided by the traditional shape descriptors.

Additionally, for both KPCA and UMAP cases, a small *p*-value produced by Friedman tests (Friedman, 1937) suggests that the three SVM classifiers, corresponding to the three sets of shape descriptors, are statistically different. Since we are comparing only three classifiers at a time, we can rely better on a Quade post-hoc pairwise test (Quade, 1979) as suggested in (Conover, 1998) (Table 3).

**Table 3:** Small Friedman and Quade post-hoc *p*-values (using *t*-distribution approximation with Bonferroni correction) suggest that different descriptors produce statistically different SVM results.

| ECT + KPCA |  |  | ECT + UMAP |  |  |
| --- | --- | --- | --- | --- | --- |
| Friedman test $p$-value: | | $1.4 \times 10^{-5}$ | Friedman test $p$-value: | | $4.4 \times 10^{-10}$ |
|  | Traditional | Topological |  | Traditional | Topological |
| Topological | $1.8 \times 10^{-11}$ | * | Topological | $8.0 \times 10^{-4}$ | * |
| Combined | $5.9 \times 10^{-5}$ | $4.4 \times 10^{-4}$ | Combined | $4.4 \times 10^{-13}$ | $7.8 \times 10^{-7}$ |

## Discussion

Traditional morphometrics has been used to reveal fundamental trends in morphological changes across space and time in ancient cereal grains (Bouby, 2001; Tanno and Willcox, 2012). Historical evidence shows that barley seeds became smaller as the crop moved from Mediterranean climates to Northwest Europe due to colder temperatures and higher sunlight variance, shedding insight on the timeline of barley domestication in Central Asia (Motuzaite Matuzeviciute et al., 2018). Similarly, grains became rounder and the spikes more compact as they moved to higher altitude sites in Nepal (Fuller and Weisskopf, 2014). Differences are more subtle if we compare accessions originating from similar regions and time periods. Geometric Morphometrics (GMM) has provided a more quantitative characterization of the grains. For example, GMM can successfully tell apart barley grains from einkorn (*Triticum monococcum*) and emmer (*Triticum dicoccum*) accessions (Bonhomme et al., 2017); it can be used to distinguish two-row vs six-row barley seeds (Ros et al., 2014); and it can establish unique morphological characteristics of land races to deduce their possible historical origins (Wallace et al., 2019).

Morphometrics has a number of drawbacks with respect to X-ray CT images. GMM requires homologous points and, although 3D and higher dimensional analysis is possible, it is usually applied to 2D images (Dryden and Mardia, 2016). Further, a geometric framework is limited to the relationship of data points to each other. We thus turn to topology. In recent years, TDA has produced promising results in diverse biological problems, like histological image analysis (Qaiser et al., 2019), viral phylogenetic trees description (Chan et al., 2013), and identification of active-binding sites in proteins (Kovacev-Nikolic et al., 2016). In plant biology, the Euler characteristic has been used successfully to define the morphospace of more than 180,000 leaves from seed plants (Li et al., 2018), and to characterize the shape of apple leaves (Migicovsky et al., 2018) and the 3D structure of grapevine inflorescences (Li et al., 2019).

The PCA of the traditional shape descriptors tends to group seeds based on accession as the largest source of variance. This observation is further supported by the Kruskal-Wallis analyses of variance (Fig. S5). The Euler characteristic however encodes additional important shape information missed by traditional descriptors. We observe that the topological shape descriptors provide better classification than the traditional shape descriptors (Table 2). Recall that we can mathematically prove that the ECT captures all the shape information, to the point that a finite topological signature can be used to reconstruct the original object (Curry et al., 2018; Fasy et al., 2019). This vast amount of information is best parsed with dimension reduction techniques, which highlight different morphology features encoded by the ECT. The biggest source of variation encoded by the ECT, rendered through KPCA, are individual panicles. This high degree of spike distinction may ignore underlying shape similarities between panicles of the same accession. In contrast, with UMAP we reduce the ECT’s dimensionality taking into account overall topology and geometry, and produce a clustering that balances both panicle-specific nuances with more general accession-based traits. This accession-vs-panicle balance is further aided by combining traditional and UMAP-reduced descriptors. In other words, the ECT is capable of capturing both panicle- and accession-specific morphological descriptors, but different dimension reduction techniques emphasize some nuances over others. The addition of traditional shape descriptors aids accession-based clustering, by supplying size-related measurements.

The majority of the accessions studied are more easily distinguished with the topological lens but not with traditional measures, with few exceptions (Fig. 7). Exceptions like Hannchen, Han River and Palmella Blue have slightly distinctive traditional trait distributions, so seed size does matter and it is important to take it into account (Fig. 5A). At the same time, we observe accessions like Alpha, Glabron, Minia, and Wisconsin Winter, that are poorly differentiated with traditional information but report considerably higher classification accuracies whenever using topological information. When looking at a more robust dimension reduction technique like UMAP, classification accuracy is increased when combined with size-related information.

**Figure 7:**
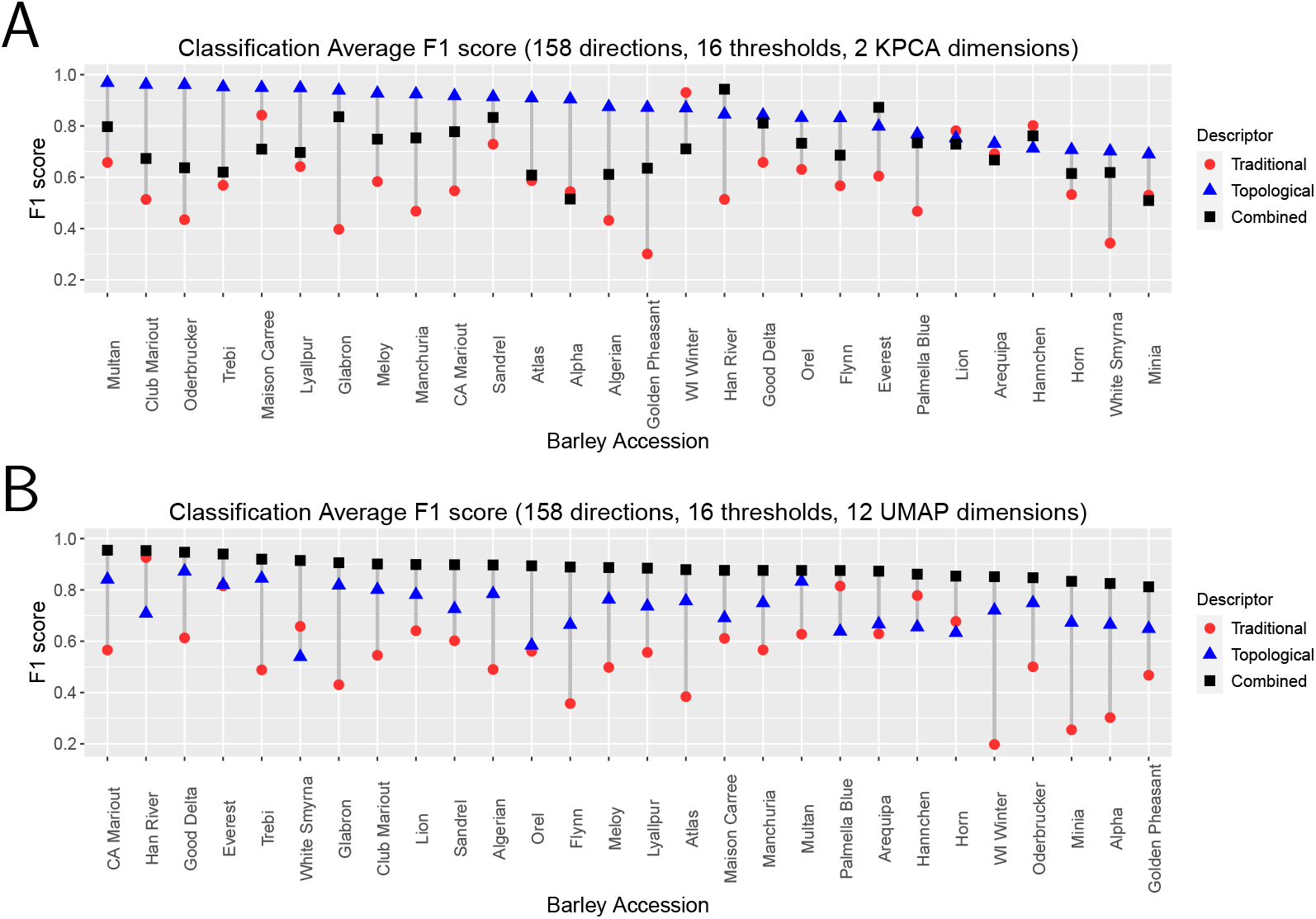
SVM classification results for individual accessions. **A.** Results when using a KPCA 2-dimension reduced topological vector. Accessions ordered according to their classification accuracy determined by the topological shape descriptors. **B.** Results when using a UMAP 12-dimension reduced topological vector. Accessions ordered according to their classification accuracy determined by the combined shape descriptors.

An exploration on the directions used to compute the ECT, reveals that the shape of the crease and bottom discriminate accessions the most (Fig. 4). These features are not directly measured with our traditional setting. By analyzing inter-vs. intra-accession variance of a large number of ECT axes and thresholds, we effectively isolate complex morphological features responsible for distinguishing selected groups. Going forward, there are a number of topics to explore with respect to the implementation of this novel approach. How are the results affected and what is the computationally feasibility, for instance, if we pick uniformly randomly distributed directions —or according to any other probability distribution— instead of polar-biased ones. Although the ECT comprehensively measures the information content of an object, different dimension reduction techniques highlight different aspects of that shape information (Fig. 6). A more systematic exploration of other dimension reduction algorithms, and classification techniques afterward is warranted moving forward.

The Euler characteristic is a simple yet powerful way to reveal features not readily visible to the naked eye. There is “hidden” morphological information that traditional and geometric morphometric methods are missing. The Euler characteristic, and Topological Data Analysis in general, can be readily computed from any given image data, which makes it a versatile tool to use in a vast number of biology-related applications. TDA provides a comprehensive framework to detect and compare morphological nuances, nuances that traditional measures fail to capture and that remain unexplored using simple geometric methods. In the specific case of barley seeds presented here, these “hidden” shape nuances provides enough information to not only characterize specific accessions, but the individual spikes from which seeds are derived. Our results suggest a new exciting path, driven by morphological information alone, to explore further the phenotype-genotype relationship.

## Software and data availability

All of our code is available at the https://github.com/amezqui3/demeter/ repository. This includes the image processing pipeline to clean the raw scans and segment the seeds (python), the computation of the ECTs (python), and the SVM classification and analysis (R). A collection of Jupyter notebook tutorials is also provided in order to ease the usage and understanding of the different components of the data processing and data analyzing pipelines.

## Acknowledgements

Daniel Chitwood is supported by the USDA National Institute of Food and Agriculture, and by Michigan State University AgBioResearch. The work of Elizabeth Munch is supported in part by the National Science Foundation through grants CCF-1907591 and CCF-2106578. Jacob Landis was supported by the NSF Plant Genome Postdoctoral Fellowship 1711807. Daniel Koenig is supported by an award from the National Science Foundation Plant Genome Research Program (IOS-2046256) and funding from the USDA NIFA (CA-R-BPS-5154-H).

## Supplement material

**Figure S1:**
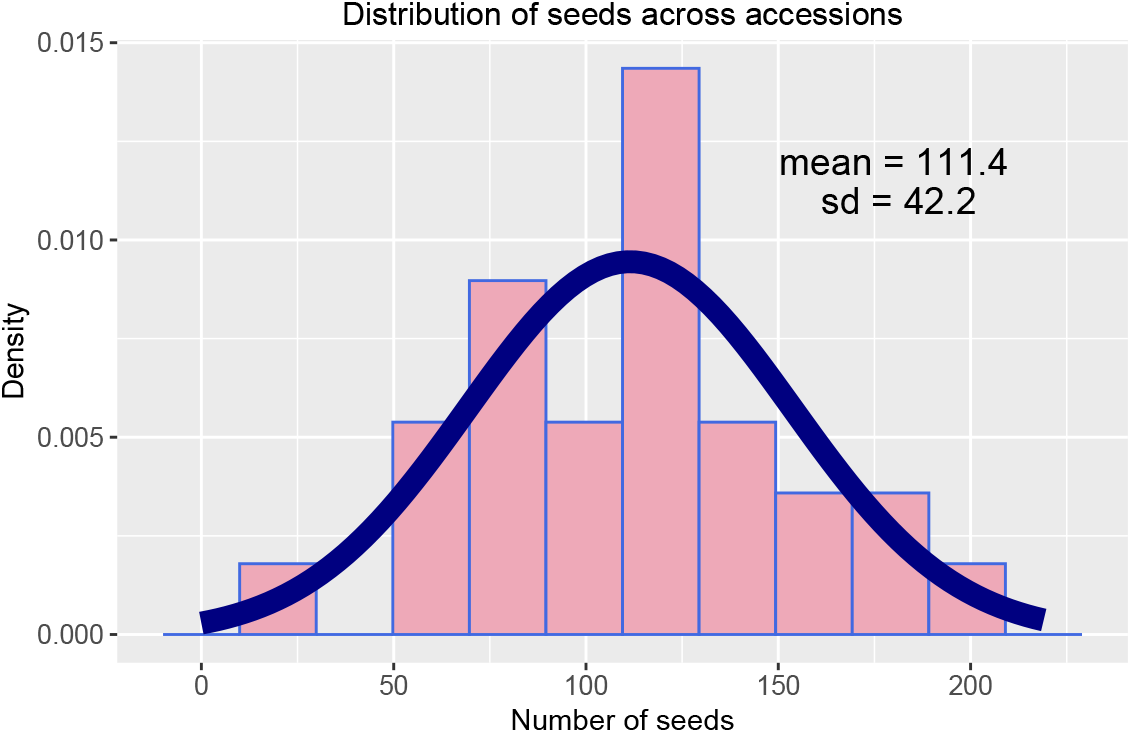
Distribution of the 3121 seeds according to their accession. The seed number values as in Table 1 have empirical mean 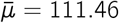 and empirical standard deviation 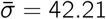. A normal distribution with these parameters is drawn on top of the histogram. Observe that all the accession seed numbers are within two standard deviations.

**Figure S2:**
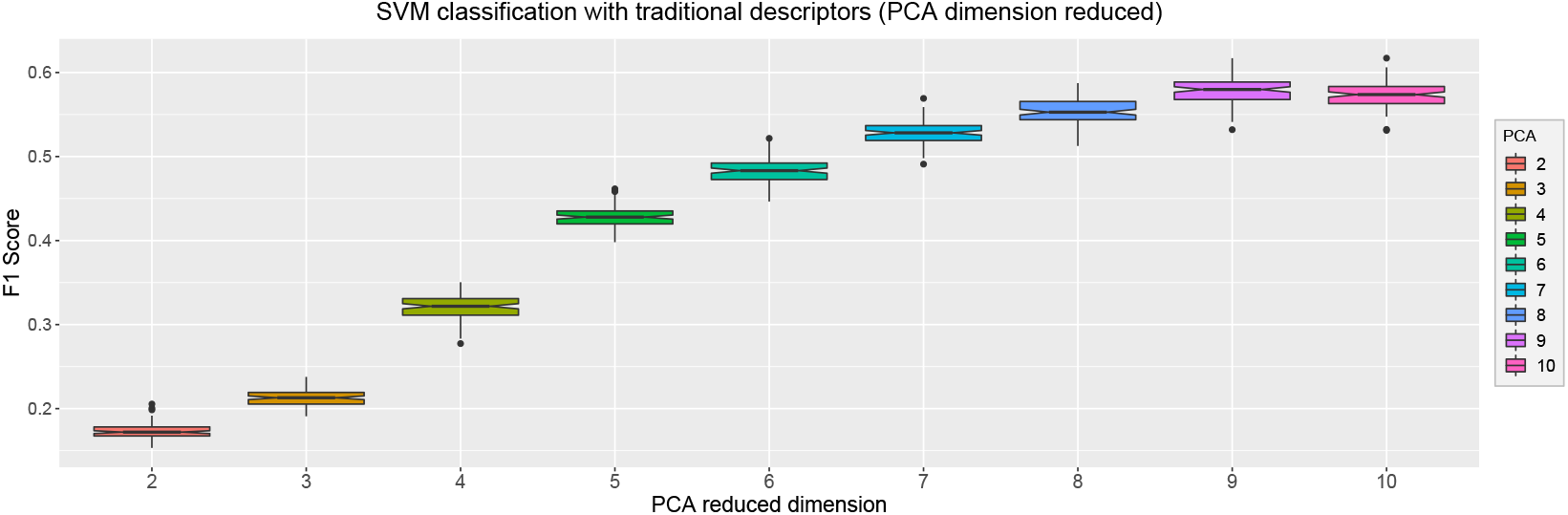
Classification results for traditional shape descriptors. After centering and scaling the traditional shape descriptors, we used PCA to reduce their dimension and then performed an SVM classification with these dimension-reduced vectors. We observe that the highest classification *F*_1_ scores correspond to the use of almost all the traditional dimensions.

**Figure S3:**
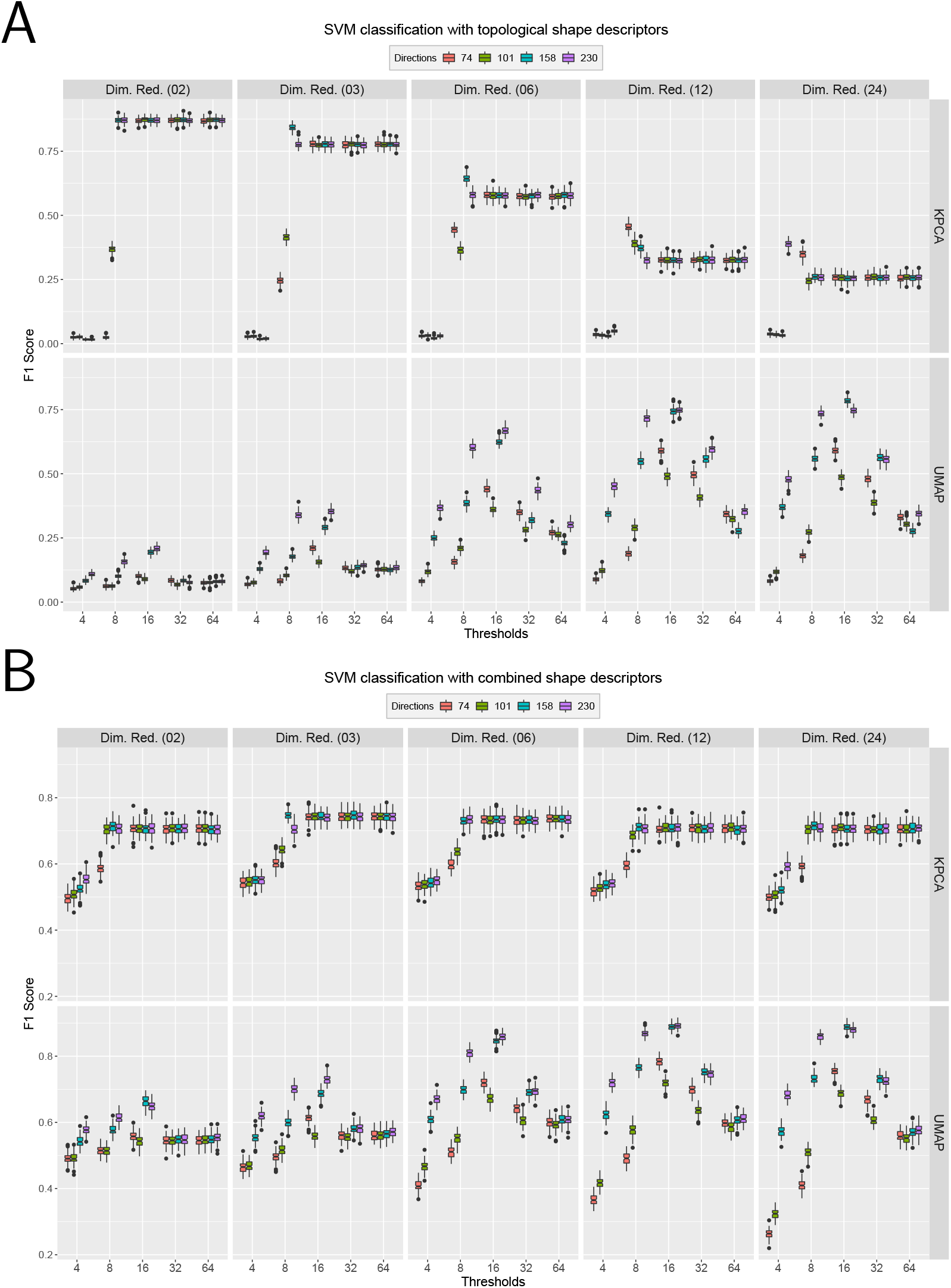
Classification results for combined and topological shape descriptors computed from different. To evaluate the ECT descriptiveness, we sought to use these ECT vectors to classify 28 different barley accessions based solely on seed morphology. The ECT was computed for different number of directions and thresholds. These high-dimensional vectors were later reduced to different number of dimensions using both KPCA and UMAP. Observe that both dimension reduction techniques summarize the ECT information in very different ways, as evidenced by the different SVM classification F1 scores when using **A.** exclusively topological information or **B.** combining both topological and traditional seed shape descriptors.

**Figure S4:**
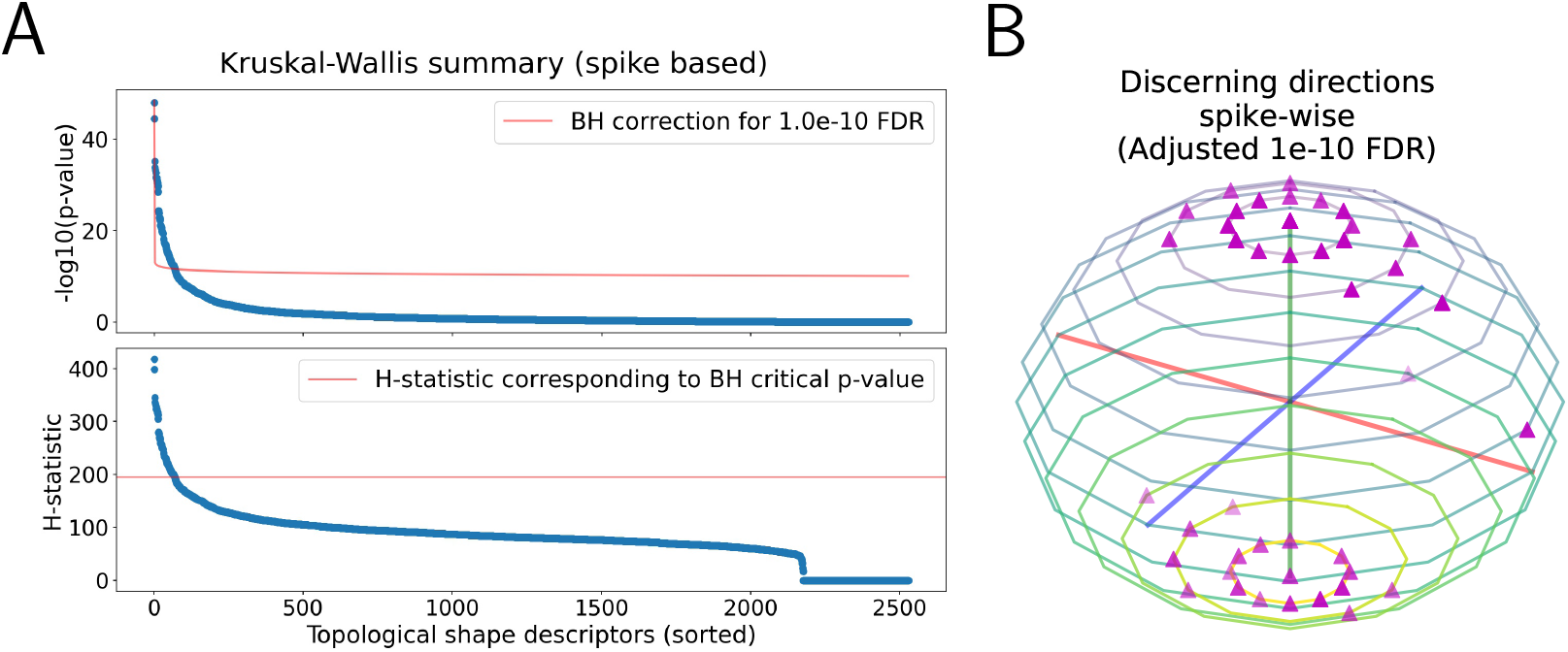
Relevant ECT directions and slices. **A.** We examine the inter-spike and intra-spike variance differences of the Euler characteristic for each direction and threshold. A Kruskal-Wallis analysis combined with a Benjamini-Hochberg multiple test correction suggests a number of discerning slices across accessions. **B.** These directions and thresholds are mostly concentrated around the poles, similar to the case of inter- and intra-accession variance case (Fig. 4).

**Figure S5:**
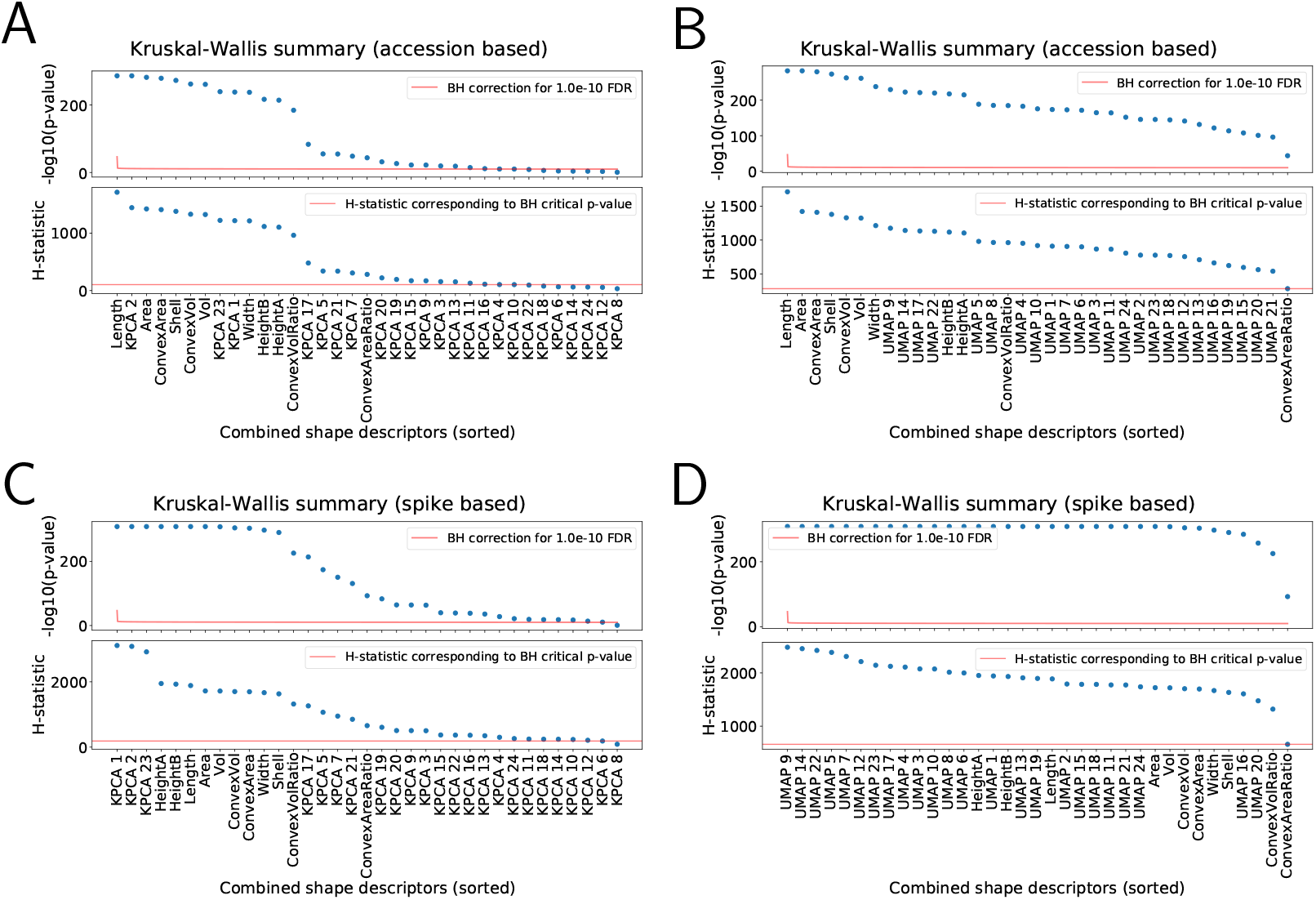
Relevant combined descriptors. Dimension-reduced topological vectors were concatenated with tradtional shape descriptors to produce combined descriptors. Kruskal-Wallis analyses reveal which descriptors explain the most inter-accession variance when the ECT was reduced in dimension with **A.** KPCA, and **B.** UMAP. Similar analyses also reveal which features contribute the most to inter-spike variance when the ECT vector was reduced with **C.** KPCA, and **D.** UMAP.

